# Attention please: modeling global and local context in glycan structure-function relationships

**DOI:** 10.1101/2021.10.15.464532

**Authors:** Bowen Dai, Daniel E Mattox, Chris Bailey-Kellogg

**Affiliations:** Department of Computer Science, Dartmouth College, Hanover, NH, USA; Program in Quantitative Biomedical Sciences, Geisel School of Medicine at Dartmouth College, Hanover, NH, USA

**Keywords:** glycobiology, glycans, lectins, deep learning, language models, attention-based learning, glycoinformatics

## Abstract

Glycans are found across the tree of life with remarkable structural diversity enabling critical contributions to diverse biological processes, ranging from facilitating host-pathogen interactions to regulating mitosis & DNA damage repair. While functional motifs within glycan structures are largely responsible for mediating interactions, the *contexts* in which the motifs are presented can drastically impact these interactions and their downstream effects. Here, we demonstrate the first deep learning method to represent both local and global context in the study of glycan structure-function relationships. Our method, glyBERT, encodes glycans with a branched biochemical language and employs an attention-based deep language model to learn biologically relevant glycan representations focused on the most important components within their global structures. Applying glyBERT to a variety of prediction tasks confirms the value of capturing rich context-dependent patterns in this attention-based model: the same monosaccharides and glycan motifs are represented differently in different contexts and thereby enable improved predictive performance relative to the previous state-of-the-art approaches. Furthermore, glyBERT supports generative exploration of context-dependent glycan structure-function space, moving from one glycan to “nearby” glycans so as to maintain or alter predicted functional properties. In a case study application to altering glycan immunogenicity, this generative process reveals the learned contextual determinants of immunogenicity while yielding both known and novel, realistic glycan structures with altered predicted immunogenicity. In summary, modeling the context dependence of glycan motifs is critical for investigating overall glycan functionality and can enable further exploration of glycan structure-function space to inform new hypotheses and synthetic efforts.

## 1 INTRODUCTION

Glycans are complex oligosaccharides often presented in branched structures attached to proteins, lipids, and RNA, with critical roles in a diverse range of biological processes [1;2]. Glycans mediate these processes through two general mechanisms [3]: (1) specific interaction and recognition of glycoforms by glycan binding proteins, such as the targeting of sialoglycans on the surface of animal cells by influenza hemagglutinin to facilitate viral entry [4], and (2) structural and biophysical effects such as the critical modulation of antibodies’ downstream effector functions by N-linked glycans [5]. Subtle modifications in glycan structures independent from functional epitopes can drastically alter glycan function, as seen in the large structural and functional shifts introduced by core fucosylation [6;7]. Additionally, lectin-glycan interactions have recently been shown to be dependent on the surrounding global structural context in which the appropriate binding epitopes are presented [8;9], e.g., as demonstrated by the glycan binding preferences of *Maackia amurensis* lectin I (MAL-I) (Fig 1, left). As glycan synthesis, purification, and presentation techniques advance and more detailed interrogations of glycans’ structure-function relationships become available [8;9], methods are needed to analyze and explore (Fig 1, middle and right) complex context-dependent aspects of glycan structure-function relationships.

**Fig 1.**
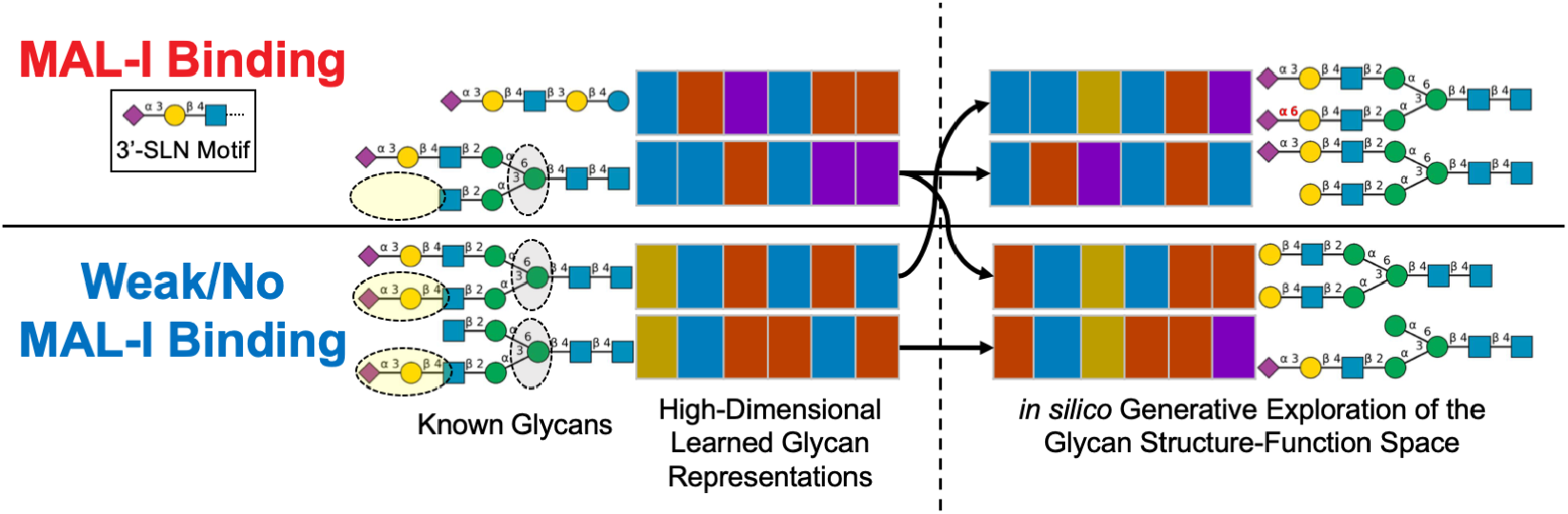
Schematic of an analytical framework to investigate context-dependent structure-function relationships in glycans. **(left)** Local context (functional glycan motifs such as 3’ sialyllactosamine (3’-SLN)) and global context (non-adjacent features indicated by dashed circles) simultaneously play critical roles in the functionalities of glycans, as demonstrated here by the glycan-binding preferences of *Maackia amurensis* lectin I (MAL-I), which recognizes 3’-SLN on linear glycans or on the *α*1,6 branch of biantennary glycans (grey circles), but only when the opposing *α*1,3 branch does not present the 3’-SLN motif (yellow circles). **(middle)** As schematically illustrated, our glyBERT approach captures such contextual relationships with high-dimensional glycan representations that can predict properties of new glycans. **(right)** GlyBERT also enables generative exploration of glycan structure-function space to provide insights into learned patterns and probe glycan diversity beyond the practical reach of synthesis, thereby generating novel hypotheses and informing experimental design.

Analyses of experimental measures of glycan functions offer a means to efficiently capture the components of diversity most relevant to functionality while also generalizing to other glycans, enabling the prediction of properties of previously untested glycans or even the generation of entirely novel glycans (Fig 1), thereby complementing and extending the impact of synthetic capabilities which remain limited relative to the scale of glycan diversity [10;11]. To date, most analytical approaches have focused on identifying common structural motifs [12] among glycans with shared properties, e.g., via subtree mining [13;14], motif enrichment analysis [15], or multiple alignment approaches [16]. These methods have been very beneficial in formalizing high level glycan-binding preferences of proteins [17]. They will continue to retain great utility due to their high interpretability, but this interpretability comes at the cost of reduced ability to capture more complex and context-dependent patterns. In general, deep learning approaches are attractive for building detailed and powerful representations from complex information [18]. The inceptive application of deep learning methodology directly to glycans, SweetTalk [19], used a “glycoword”-based language model to successfully demonstrate that deep modeling could capture evolutionary and functional patterns in glycan structures [20]. The follow-up SweetNet approach [21] obtained increased prediction performance by utilizing graph-based glycan encodings and convolutional neural networks, thereby addressing SweetTalk’s failure to capture the branched nature of glycan structures. While convolutional approaches can capture local context, they do not preserve global relationships [22] which are critical in glycan functionality [8;9]. Another very recent effort, GlyNet [23], implemented a fingerprinting approach to predict relative strengths of protein-glycan interactions, but faces similar limitations as SweetTalk & SweetNet in terms of disregarding global structural context.

In order to capture local and global context-dependent patterns driving glycan structure-function relationships, we introduce here a new method based on the powerful deep language model known as BERT (Bidirectional Encoder Representations from Transformers) [24]. Our implementation, called glyBERT, utilizes a flexible glycan encoding based on a branched biochemical language, and presents encoded glycans to an attention-based transformer network architecture [22], thereby learning to pay attention to particular local patterns that are important within particular global contexts. GlyBERT effectively learns glycan representations that capture functional pieces of glycan structural diversity in their local contexts while also incorporating the critically important global contexts of entire structures. This enables the model to obtain state-of-the-art performance in predicting properties of other glycans. Furthermore, glyBERT supports a novel proof-of-principle generative process to explore glycan structure-function space, providing insights into what the model has learned as determinants of different glycan properties, as well as potentially informing experimental design and focusing *in vitro* synthesis efforts [25].

## 2 RESULTS

GlyBERT was trained on 80% of a set of 16,048 glycans from the SugarBase database [26], labeled with glycoprotein linkage, immunogenicity, and taxonomic origin. The glycans from the withheld 20% were used to explore the context-dependent representations learned by glyBERT relevant to these properties (subsection 2.1) and to quantitatively evaluate prediction performance (subsection 2.2). GlyBERT was further trained and evaluated for prediction of lectin-glycan recognition using glycan microarray data for 20 separate lectins (subsection 2.2). Finally, glyBERT was leveraged to generatively explore glycan structure-immunogenicity space in a case study application of our new algorithm for generative optimization of glycan structure-function relationships (subsection 2.3).

### 2.1 Attention-based modeling with glyBERT learns representations capturing effects of local and global structural context

GlyBERT, like all machine learning methods, essentially transforms its training data into a novel representation (schematically illustrated in Fig 1), enabling it to generalize to new instances (here glycans) based on how their representations compare to the training examples. In order to make high-quality predictions about properties of the new glycans, it is critical that the learned representations suitably capture important, generalized components of glycan structures. We explored the extent to which the glycan representations that glyBERT learned from the training set generalized to new glycans in the held-out set, first in terms of global structural contexts of whole glycans, and then in structural contexts of constituent monosaccharides. Since the learned representations within the deep learning model are hard to interpret on their own, they were transformed and visualized via Uniform Manifold Approximation and Projection (UMAP) dimensionality reduction [27].

GlyBERT learned glycan representations that effectively separated out the effects of local structural motifs based on global structural contexts, differentiating glycan structures on the basis of immunogenicity, glycoprotein linkage, and taxonomic origin (Fig 2A). The immunogenicity of the same immunogenic motifs was modulated by global structural context captured in the glyBERT-learned representations: Lewis-Y antigen-presenting immunogenic glycan SBID3980 was grouped closer to other immunogenic glycans while the exact same structure with the addition of a reducing-end terminal *β* -linked glucose (non-immunogenic glycan SBID4613) was grouped separately with other non-immunogenic glycans. Similarly, the immunogenic I antigen (SBID5978) is no longer immunogenic when presented on an O-GalNAc glycan where either the reducing-end terminal GlcNAc is substituted for a GalNAc (SBID2468) or a GalNAc residue is added to the reducing end (SBID12833), with clear separation between these glycan representations matching their immunogenic states based on the global context of the I antigen presentation, despite high similarity between the structures.

**Fig 2.**
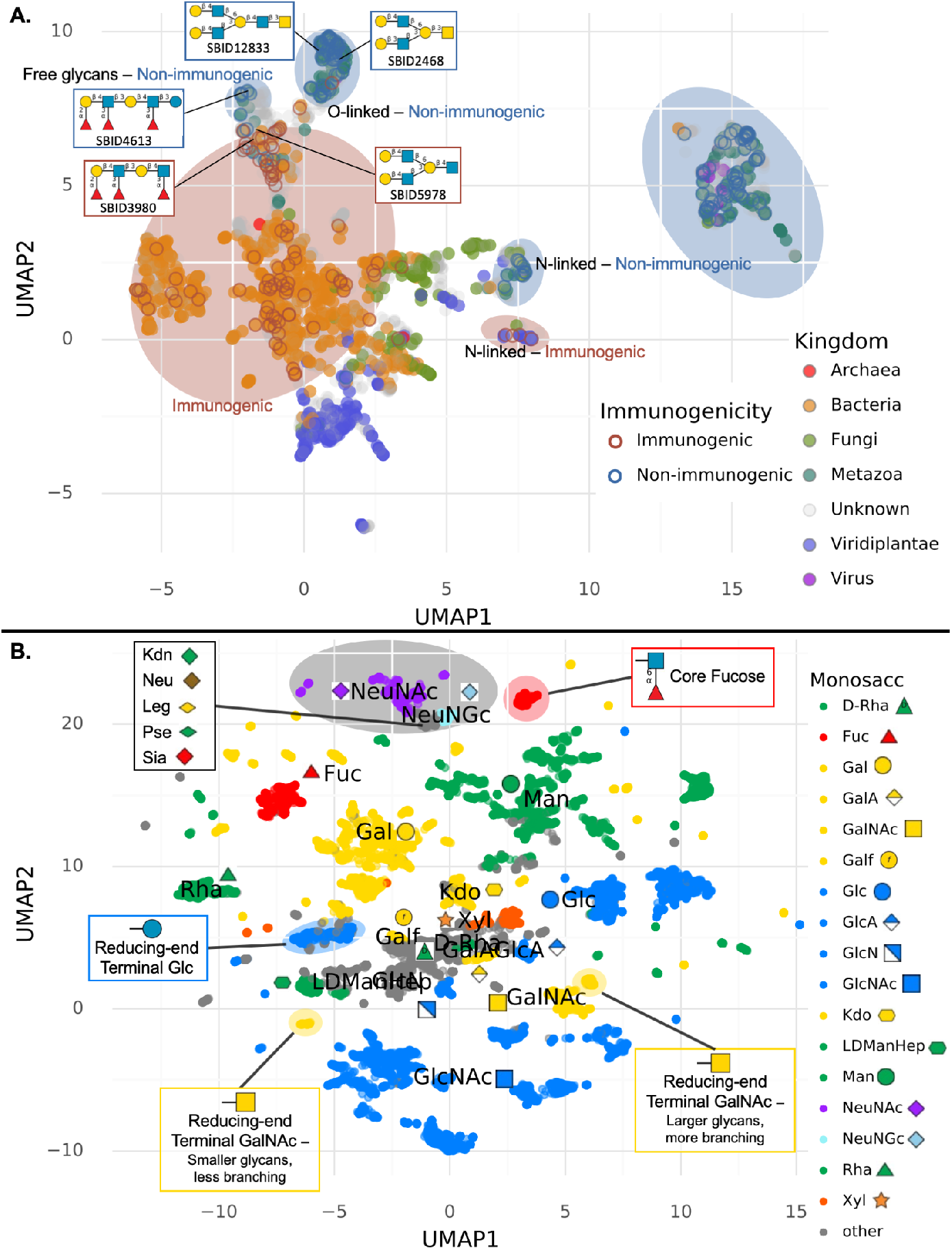
GlyBERT-learned glycan representations capture global and local contextual relationships that are consistently manifested in an independent validation set of glycan structures. **(A)** This UMAP visualization places withheld glycans with similar glyBERT representations close to each other, colored by taxonomic origin at the kingdom rank and outlined by immunogenicity. Clusters of glycans are labeled and circled, showing that glyBERT grouped the previously unseen glycans by shared immunogenicity and/or glycoprotein linkage. For further interpretation, some example glycan structures from both training and testing tests are provided, illustrating that glyBERT’s representations for otherwise very similar glycans were shifted based on the influence of global context on immunogenicity (indicated by colored boxes); e.g., non-immunogenic SBID4613 and immunogenic SBID3980 both display the Lewis-Y antigen motif but in different contexts, and immunogenic SBID5978 and non-immunogenic SBID2468 & SBID12833 display the I antigen motif in different contexts. **(B)** This UMAP visualization places individual constituent monosaccharides of withheld glycans close to other monosaccharides with similar glyBERT representations, with the most common ones colored and labeled. It is apparent that the same monosaccharides are represented similarly unless their local or global structural contexts differ. GlyBERT represented fucose in a core fucosylation local context (circled in red) differently from other fucose residues, as it did for glucose in a reducing-end terminal local context (circled in blue). O-linked reducing-end terminal GalNAc residues formed two distinct clusters (circled in yellow), separated by the size and degree of branching of the corresponding glycans (global context). Additionally, glyBERT grouped together the distinct but chemically similar nonulosonic acid monosaccharides (grey circle) based on similar contextual presentations.

In addition to evaluating similarities of entire glycans, constituent monosaccharides can be compared based on their individual “contributions” to the overall glycan representations. The constituent monosaccharide representations still capture their local and global context due to the attention-based architecture of glyBERT. We see that the same monosaccharide components of the withheld glycans generally shared similar representations (Fig 2B), although glyBERT representations also captured dramatic differences in the local glycan context in which they appeared. This can be seen in the distinct groupings of fucose saccharide residues in a core fucosylation context from the rest of the fucose residues, as well as the separation of glucose as a reducing-end terminal saccharide from the rest of the glucose residues. These local contexts are clearly represented as separate from other occurrences of the same monosaccharides, as would be expected from a good representation, since reducing-end terminal glucose is sufficient to mitigate the immunogenicity of a Lewis-Y antigen-presenting glycan (Fig 2A) and core fucosylation can induce significant conformational and functional changes in glycans [6;7]. Learned monosaccharide representations also captured differences in the global context of their corresponding glycan structures, seen in the two separate groups of reducing-end terminal O-linked GalNAc residues distinct from other GalNAc residues based on distinct local contexts and separated from each other based on global structural contexts depending on the size and degree of branching of the entire glycan. On the flip side, different and separate monosaccharides that appear in very similar global and local contexts had similar representations as learned by glyBERT, indicating generalizabilty to novel glycan structures. This can be seen in the grouping composed exclusively of nonulosonic acid monosaccharides, the larger class containing sialic acid monosaccharides and other negatively charged 9-carbon monosaccharides [28]. Notably, representations of the related 8-carbon Kdo monosaccharide grouped separately from the nonulosonic acid monosaccharides despite shared functional groups and biosynthetic pathways [28].

### 2.2 Contextual glycan representations learned by glyBERT enable state-of-the-art predictive performance

As a demonstration of the utility of these learned context-dependent glycan representations, glyBERT’s ability to predict glycoprotein linkage state, immunogenicity, and taxonomic origin for the glycans from the withheld 20% from SugarBase was compared to that of three recent deep learning-based approaches: SweetTalk [19;20], SweetNet [21], and GlyNet [23]. All models predicting glycoprotein linkage performed very well, with glyBERT and SweetTalk achieving 99% accuracy, suggesting that this structural association is fairly easily represented. However, for immunogenicity, glyBERT (98.0% accuracy) substantially improved on SweetNet (94.6%) and GlyNet (95.4%), which had improved on the first-generation SweetTalk (91.7%), indicating benefit from explicitly accounting for branched structures (as done by SweetNet, GlyNet, and glyBERT) and further benefit provided by the use of attention-based modeling to capture context (glyBERT). For taxonomic origin classification (Fig 3A), glyBERT displayed the highest accuracy at each rank, with greater increases in performance compared to SweetNet at the phylum, class, and order taxonomic ranks.

**Fig 3.**
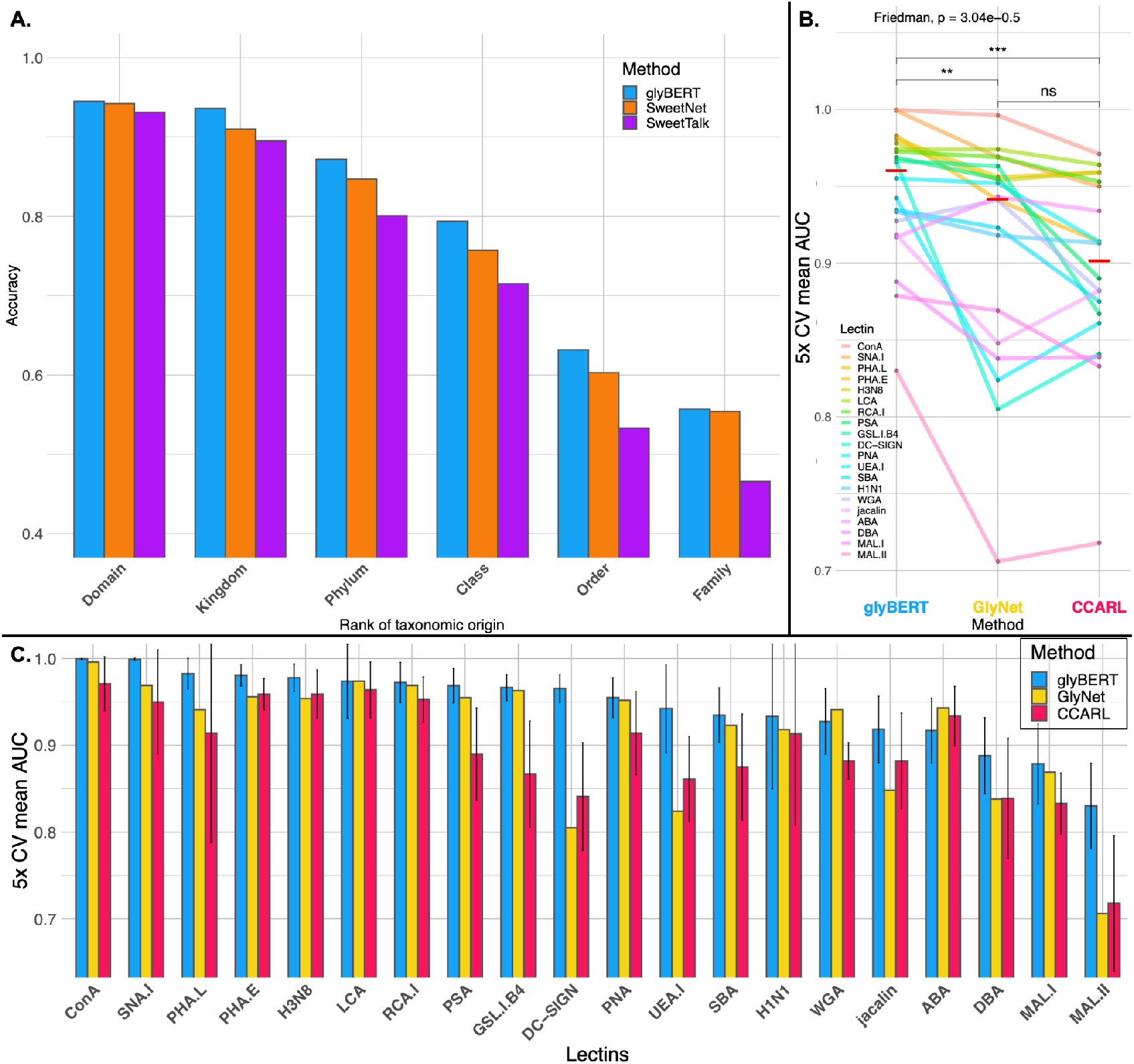
Global & local glycan structural context in learned representations enable state-of-the-art prediction of glycan properties and lectin-glycan interactions. **(A)** GlyBERT had higher taxonomic classification accuracy than SweetTalk and SweetNet at each taxonomic rank. **(B & C)** GlyBERT demonstrated significantly improved predictive modeling of 20 lectins’ glycan-binding preferences when compared to GlyNet and CCARL as seen in the parallel coordinates plot with median performances indicated by red bars (B) and in the bar plots with error bars indicating standard deviation across the 5-fold cross-validation (C). Significance was measured via a Friedman test followed by *post hoc* pairwise Wilcoxon signed-rank tests and Holm-Bonferroni correction (*: p_adj_ *<* 0.05, **: p_adj_ *<* 0.01, ***: p_adj_ *<* 0.001: ***).

To further evaluate the benefit of attention-based glycan modeling for complex structure-function relationships, the prediction of functionally significant lectin-glycan interactions was considered, since recent studies have shown the glycan-binding preferences of lectins to be more context-dependent than previously understood [8;9]. GlyBERT was compared to GlyNet [23] and a recent, well-performing motif-centric subtree mining approach known as the Carbohydrate Classification Accounting for Restricted Linkages (CCARL) method [14]. All methods were trained and evaluated on the same set of 20 lectins with the same lectin-glycan interaction labels determined from microarray data from the Consortium for Functional Glycomics (CFG) [29] identically processed and divided into folds for training and cross-validation (CV) as provided by Coff et al. [14]. The glyBERT model trained above for glycan properties provided the basis for fine-tuning on the CFG microarray data, resulting in 20 separate models for the 20 lectins. Results for the other methods were previously reported [14;23].

Performance was measured via area under the receiver operating characteristic curve (AUC), yielding a median AUC over the 20 lectins of 0.960 for glyBERT, demonstrating an increase in overall performance accuracy compared to GlyNet (0.946) and CCARL (0.896) with a statistically significant difference in ranked performance on matched lectins between glyBERT and the other two methods (Fig 3B). The greatest increases in performance were seen for *Maackia amurensis* lectin II (MAL-II) and dendritic cell receptor DC-SIGN (Fig 3C), indicating glycans on the CFG microarray likely manifested contextual dependencies. Indeed, DC-SIGN is known to have dual specificities for oligomannose glycans and blood group epitopes [30], with contextual oligomannose-preferences dependent on both the branch containing its primary motif and the mannose-content of surrounding branches [9]; furthermore, MAL-II has similar glycan preferences as MAL-I (Fig 1) but preferentially recognizes certain sialylated glycans when they are O-linked [31].

### 2.3 Deep generative exploration of the glycan structure-function space reveals contextual features discriminating immunogenic glycans

As illustrated in the preceding sections, machine learning models like glyBERT and others generalize structure-function relationships learned from known glycans in order to make predictions for new glycans. While to our knowledge no previous work has gone further, such models also may also be used to *generate* new glycans according to unwritten specifications of desired structure-function relationships. This provides the opportunity to probe the vast diversity of glycan structures, aiding in hypothesis generation and informing directions for glycan-synthesis efforts. Additionally, it allows for *post hoc* analysis of captured patterns that could be distilled into more interpretable summarizations, e.g., by MotifFinder [15].

While full validation of computationally-generated glycans would require experimentation beyond the scope of the present study, we demonstrate here proof of concept by working with a subset of glycans that are structurally related but display different immunogenicity. We start with non-immunogenic glycans and apply a generative optimization approach, substituting monosaccharides at each step so as to push the glycans toward being immunogenic, based on probability gradients according to the glyBERT model. This enables an exploration of the glycan space near the starting non-immunogenic glycans, leading to some of those known be immunogenic along with novel glycans predicted to be immunogenic due to glyBERT’s contextual assessment of their motifs. This process reached the known immunogenic counterparts within 2-3 steps (Fig 4A) for four of the eight starting non-immunogenic glycans for which known immunogenic counterparts exist. For three of remaining four it arrived upon known, unlabeled glycans that exactly matched existing immunogenic glycans other than the anomeric configuration of the glycosidic bonds in the complete set of generative paths (Fig S2). In such a generative exploration, glycan structures that are generated more frequently (node size) might be more realistic (and here, more immunogenic) than others; notably the most frequently generated immunogenic structures in this case study were in fact known glycans. Generative steps that reverse direction to go against the probability gradient might indicate regions with few “nearby” realistic structures, as is likely the case for the non-realistic dead-end path from NeuAc(*α*2-3)Gal(*β* 1-3)GalNAc to GalNAc(*α*2-3)Gal(*β* 1-3)GalNAc.

**Fig 4.**
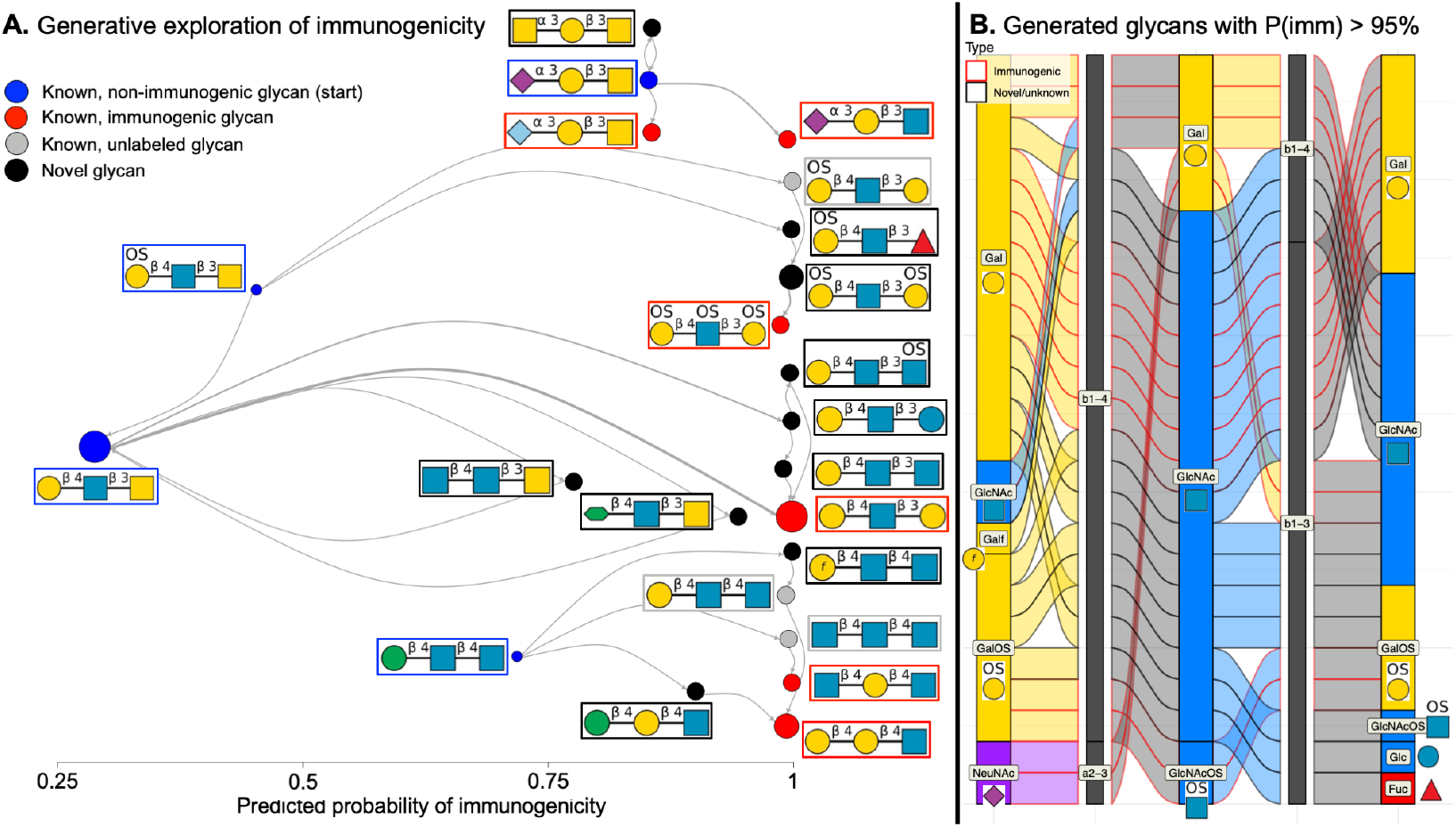
Generative exploration of glycan structure-immunogenicity space. **(A)** Generative optimization starting from non-immunogenic glycans (blue) introduced monosaccharide substitutions to increase the probability of immunogenicity (x axis), allowing for the exploration of “nearby” known and unknown glycan structures (grey & black respectively), with some known glycans also labeled as immunogenic (red). Node sizes correspond to the number of times a glycan structure was reached along any of the generative paths, and edge widths correspond to the number of times a step between two glycan structures occurred across all the generative processes. Known immunogenic glycan structures were recovered for half of the starting, non-immunogenic glycans in only a few generative steps, demonstrating the process is able to generate realistic glycan structures and can optimize the generated structures for properties of interest. **(B)** This case study revealed features of glycans determined by glyBERT to confer immunogenicity in this structural context as shown in this alluvial plot [32] summarizing the structures of generated glycans with predicted probability exceeding 95%, including Gal and Gal derivatives in the non-reducing-end terminal position, GlcNAc in the middle position, and GlcNAc or Gal in the reducing-end terminal position.

Features in glycan structures deemed by glyBERT to be critical for immunogenicity are revealed by the generated glycans with the highest predicted probabilities of immunogenicity (Fig 4B). While Gal and Gal derivatives at the non-reducing-end terminal position and GlcNAc monosaccharides at the middle position generally dominate the generated glycans predicted to be immunogenic, the two non-immunogenic glycans with the lowest probability of immunogenicity share this monosaccharide composition at these positions, indicating the learned significance of GlcNAc and Gal monosaccharides but specifically not GalNAc at the reducing-end terminal position in this context. The substitution of NeuAc for the generally immunogenic NeuGc monosaccharide [33] imparted less gain in predicted immunogenicity than substituting the reducing-end terminal GalNAc for GlcNAc in the generative paths shown at the top of Fig 4A. These trends were generally recapitulated in the complete set of generative paths (Fig S2).

## 3 DISCUSSION

As experimental techniques evolve to enable richer studies of complex glycan structures, analytical techniques to study structure-function relationships must also evolve to effectively capture complex and context-dependent associations between glycan properties and the global contexts in which functional motifs and epitopes are presented. To suitably learn and leverage context-dependent glycan structure-function relationships, we developed an attention-based model (glyBERT) using minimally processed glycan encodings. Building on the pioneering efforts of Bojar et al. [20] and Burkholz et al. [21], we further demonstrated the value of deep learning in glycobiology, here by representing, leveraging, and revealing contextual glycan features. We showed that the learned glycan representations are sensitive to both local glycan features and global contexts in relation to a variety of glycan functions and labels, that they support state-of-the-art predictive performance of glycan properties from structures, and that they enable exploration of structure-function relationships while generating novel glycans.

One limitation of the glyBERT approach was the lack of positional information for monosaccharide modifications in the training data from the SugarBase database; this limitation manifested for example in the relatively poor prediction performance for lectin ABA in Fig 3C & Table S1. In general, distinguishing monosaccharides and modifications would allow the model to more easily learn relationships between rare monosaccharides and rare modifications using independent occurrences instead of relying on co-occurrence. And while glyBERT learned implicit relationships between chemically similar monosaccharides (Fig 2B), additional pre-training based on stereochemical structure and composition of monosaccharides and their modifications could further improve generalizability to more rare residues.

While the generative case study presented here is limited in scope due to our desire to recapitulate some known, closely-related immunogenic/non-immunogenic glycan pairs, the same framework could be applied to any property of interest and the gradient-based sampling method could be expanded to explore much more drastic changes to glycan structures. These expansions would truly enable deep glycan structure-function space exploration and contribute contextual-dependencies to glycoengineering efforts that might be applied to the design of competitive inhibitors or glycomemetics.

## ACKNOWLEDGMENTS

We would like to thank Zach Klamer for discussion while the manuscript was being prepared, as well as Professors Margaret Ackerman, Karl Griswold, and Jiwon Lee of the Dartmouth Thayer School of Engineering, and their lab members, for their continued feedback on this project and many others at all stages of development.

## AUTHOR CONTRIBUTIONS

BD Conceptualization (Equal), Writing – Original Draft Preparation (Equal), Writing – Review & Editing (Equal), Methodology (Lead), Software (Equal), Investigation (Equal), Formal Analysis (Equal), Visualization (Equal)

DEM Conceptualization (Equal), Writing – Original Draft Preparation (Equal), Writing – Review & Editing (Equal), Methodology (Equal), Software (Equal), Investigation (Equal), Formal Analysis (Equal), Visualization (Lead), Data Curation (Lead), Funding Acquisition (Support)

CBK Conceptualization (Equal), Writing – Original Draft Preparation (Supporting), Writing – Review & Editing (Equal), Methodology (Supporting), Project administration (Lead), Visualization (Equal), Funding Acquisition (Lead)

## DATA AND CODE AVAILABILITY

The glycan structures and associated labels used in this study were retrieved from SugarBase v2.0 (https://webapps.wyss.harvard.edu/sugarbase/) [26]. Lectin-glycan interaction data were retrieved from the supplemental information of Coff et al. [14]. All code and data used to perform this analysis, including training/testing splits and class labels, is available at https://github.com/demattox/glyBERT.

## 4 METHODS AND MATERIALS

A total of 16,048 unique glycan structures were encoded to an array-format compatible with glyBERT while maintaining all relevant chemical and branching information (Fig S1A, subsection 4.3). These glycans were split into a training and test set following the same 80*/*20 split strategy as SweetNet [21] although the exact split used in that study was not available for direct comparison. Within the training set, glyBERT was pre-trained by predicting masked pieces of these structures as well as relevant labels (Fig S1B, subsection 4.5). For each prediction task with reported performance, the pre-trained glyBERT model was fine-tuned by training specifically for each individual task, allowing the model to retain the most relevant information learned in the pre-training while focusing more specifically on each different task separately (Fig S1C, subsection 4.6). Generative exploration was performed by using the immunogenicity prediction layer to calculate probability gradients and substitute individual monosaccharides that would lead to glycan structures with higher overall predicted probability of immunogenicity (Fig S1D, subsection 4.7).

### 4.1 Data collection and cleaning – SugarBase glycans

From the 19,299 available glycan entries in the SugarBase v2 database [26] downloaded on November 11^th^ 2020, 1,145 entries were eliminated because the glycan IUPAC was improperly or unexpectedly formatted. This included ambiguous or unmatched parentheses/brackets, incomplete orphaned glycosidic bonds, and branching brackets around the first monosaccharide or group of monosaccharides in the IUPAC, indicating that a portion of the glycan might be missing (e.g., SBID 712: **[Glc(a1-6)]4)**Glc(a1-4)Glc(a1-4) … Glc(a1-4)Glc). An additional 1,319 entries were eliminated after discovering they were duplicates of glycans stored in other entries in SugarBase with the order of the branches rearranged in their IUPAC representations (e.g., from SBID8088 & SBID13110: GalNAc(b1-4)[NeuNAc(a2-3)]Gal & NeuNAc(a2-3)[GalNAc(b1-4)]Gal). To facilitate systematic representation of branching, one glycan was omitted as an outlier with 25 branches since the next highest observed number of branches in a glycan was 11. Lastly, 786 glycans were omitted with any of the 531 very rare monosaccharides occurring in fewer than four glycans as there were not enough examples of these monosaccharides to allow for sufficient pre-training. Of the 19,299 original entries, we used the 16,048 unique glycans with sufficiently common monosaccharide vocabularies for the remainder of this work.

Taxonomy labels at varied taxonomic rank were recovered from the provided lists of species of origin for each SugarBase glycan entry by searching for the provided genus-level taxonomic ID (taxid) in the Python implementation of the NCBI’s taxonomy tool Ete3 v3.1.2 [34] (local data written on December 29, 2020). Sugarbase provided 835 unique genera, and all but 15 were successfully matched to NCBI taxids. Of the 15 missing genera, 11 were viruses without scientific names; these were manually added to the “Virus” superkingdom/domain. The remaining 4 were from a typo or cases of deprecated or unofficial genus classifications and were manually corrected (*Columbia livia* → *Columba livia, Kinetoplastids* → *Kinetoplastida, Actinogyra* → *Umbilicaria, Arecastrum* → *Syagrus*). Some species/genera did not have defined taxids at certain taxonomic ranks. As taxonomic origin was not the primary goal of our study, this was manually addressed at the kingdom level only since NCBI taxonomy only categorizes eukaryota at the kingdom level and there were the most missing labels at this rank. To recover some labels for a significant portion of the glycans at this rank, “Bacteria”, “Virus”, & “Archaea” labels were carried over from the superkingdom/domain rank to the kingdom rank. Beyond the aforementioned adjustments, only the NCBI defined and provided taxids were used in all other cases to ensure the quality and reproduciblity of labels used to train the model.

Labels were rectified over replicated glycans as follows. For linkage or immunogenicity labels from replicated glycans, the label was set to “None”/”Unknown” when directly conflicting labels were present, or the definitive label was used if other replicates for a glycan were unlabelled. For species of origin labels from replicated glycans, the set of all observed species of origin from all replicates of the glycan was used.

### 4.2 Data collection and cleaning – CFG glycans

Unique glycans from the Consortium for Functional Glycomics (CFG) microarray v5.0 were processed to match the specific IUPAC formatting utilized by SugarBase. To ensure patterns and relationships learned from training on SugarBase glycans could be leveraged, monosaccharide names from the CFG glycans were manually adjusted to match the names used in SugarBase glycans, including the reordering of modifications (GlcNA → GlcAN) and the removal of position specific information from sulfation modifications ((3S)Gal / (6S)Gal / (3S)(6S)Gal → GalOS).

Lectin binding to CFG microarray glycans was determined from data provided in supplemental file 6 of Coff et al. [14], splitting binding interactions into the bound class or the unbound class based on the Median Absolute Deviation (MAD) technique, while discarding intermediate interactions to maximize type I and type II error control.

### 4.3 Glycan encoding

Glycan IUPAC strings were processed into custom tree objects built from anytree v2.8.0 [35], placing each monosaccharide into a node while recording glycosidic linkage information and anomeric conformation (Fig S1A). A “START” token was placed before the reducing-end terminal monosaccharide (root node) and “END” tokens were added after each non-reducing-end terminal monosaccharide. The “START” token allowed for embedded representations of entire whole glycans and the “END” token was designed to allow for more dramatic changes to the glycan structure in future generative processes. These tree structures were then encoded into seven-column arrays with each node placed into its own row, recording the following information for each saccharide in its columns: (1) identity, represented by a single number corresponding to the rank order of the unique monosaccharides by frequency; (2) anomeric conformation, *α, β*, or Unknown; (3) indices of carbons involved in glycosidic bonds to other monosaccharides; (4) position within the branched structure glycan in a “subway line” approach described in the following paragraph; (5) index of the carbon linking to the parent monosaccharide; (6) depth, or distance from the root node; (7) index in a list of monosaccharides at the same depth on different branches, ordered by following the lowest carbon number at each branch point in a depth-first tree traversal. The carbons involved in glycosidic bonds to other monosaccharides were stored as 9-bit binary numbers, with each position representing a carbon number for a given monosaccharide participating in a glycosidic bond set to 1 and the remaining positions set to 0.

Glycan branching was explicitly accounted for in this encoding with the “subway line” approach illustrated in Fig S1A, where lines were drawn along monosaccharide nodes from each non-reducing-end terminal monosaccharide to the reducing-end terminal monosaccharide (root node) resulting in what appears like a map of subway lines that account for each branching event in a glycan. For each monosaccharide, the branches (or subway lines) that pass through that node (or subway station) were recorded in an 11-bit binary number, with the bits corresponding to branches passing through the node set to 1. An 11-bit binary number was used to have a consistent branching representation for all glycans with up to 11 branches. The 9-bit glycosidic bond binaries and 11-bit branch position binary numbers were converted into decimal numbers in the final encoding to simplify representation and reduce dimensionality. Depicted glycan representations follow the SNFG system [36] and were rendered using DrawGlycan-SNFG [37].

### 4.4 Architecture and Implementation

GlyBERT was implemented in Pytorch [38] & Fairseq [39] following the architecture diagrammed in Fig S1B. For embedding layers, the original BERT architecture embedded all different input feature encodings to the same dimension and combined them, adding positional (index of word) and segment (context information) embeddings to token embeddings. To adapt this approach to glycans, the monosaccharide encodings described above (anomeric state, carbon linkage, branch, parent, and depth) were embedded and added to the monosaccharide embedding. Learnable embedding layers were utilized for these different types of encoding features. For the transformer architecture, we used the same 12 multi-head self-attention layers (each with 12 heads) as the BERT_base_ model. The classifier layers were composed of 6 Multilayer Perceptrons (MLPs) with one for each of our customized pre-training tasks. GlyBERT was optimized with Adam [40].

### 4.5 Pre-training

In order to begin building representations of the glycans that capture the structural diversity of all available glycans from SugarBase with contextual relevance to biologically relevant properties, we employed a pre-training process inspired by the original BERT work [24]. We used six pre-training tasks, some semi-supervised and some supervised:

#### Semi-supervised

To build an implicit understanding of the different monosaccharides and their glycosidic bonds and their use in glycans in different contexts, pieces of each glycan structure (monosaccharide identity, anomeric conformation, and glycosidic bond information) from the the training set were masked, and the model was trained to predict the masked information based on its surrounding local and global context in the glycan structure. This approach treats monosaccharide types as words connected in branched sentences and follows pre-training strategies of conventional language models. All three masked prediction tasks followed the “mask LM” (MLM) procedure [41], here masking 25% of each glycan input (before padding) independently by a “MASK” tag. Only the cross entropy loss of masked tokens was computed, following Devlin et al. [24].

#### Supervised

To incorporate biologically relevant information into the glycan representations, supervised tasks included predicting N/O glycoprotein linkage, immunogenicity, and taxonomic origin. MLPs for these tasks took the embeddings of the “START” tokens as representative inputs for the whole glycans. Cross entropy loss was used for N/O linkage and immunogenicity prediction. As glycans can be found in multiple organisms, taxonomic origin prediction was treated as a multi-class, multi-label prediction task and binary cross-entropy was employed.

### 4.6 Fine-tuning

GlyBERT was fine-tuned separately to study each specific glycan property for which predictive performance was reported (Fig S1C). In this step the other MLPs’ prediction heads were simply disabled.

For the downstream fine-tuning task of predicting binding information on CFG data, we used the pre-trained model without its classification layers. A 3-layer MLP, taking the “START” token embedding as input, was then appended to the pre-trained model. Since one model was trained for each lectin’s binding activity, we used binary cross entropy loss. Models were trained using the same stratified 5-fold cross-validation as GlyNet and CCARL and optimized with Adam [40].

### 4.7 Generative optimization

To optimize glycans for a given property, two functions are needed: a differentiable function that is able to evaluate how well the current glycan meets the desired property and a function to map glycan representations in the latent space back to the encoding space [42]. In the glyBERT architecture, the prediction layer serves as the property evaluation function; since it is a MLP that combines a sequence of linear operations and differentiable non-linear activation functions, it is able to predict the property of interest and is differentiable. The MLM layer serves as the function to map a glycan’s latent space representation back to a probability distribution belonging to the encoding space.

With these two functions in hand, the pipeline for generating glycans follows naturally (Fig S1D), based on the embedding, attention, MLM, and prediction layers from the trained glyBERT model. For a given glycan structure used as the starting point, its encoding was input into the embedding layer, returning an initial embedding. The multi-head attention layer provided the glycan representation and monosaccharide token embeddings in the latent space. Monosaccharide embeddings were used to compute the original monosaccharide type probability distribution, *M*_0_, by the MLM layer. The glycan representation was then passed through the immunogenicity label prediction layer (serving as the property evaluation function) and a cross entropy loss against the positive (immunogenic) label was computed.

To update the embedding in the latent space, the network parameters were frozen and the gradient of loss with respect to the embedding was calculated. The embedding was then updated by the gradient with a learning rate of 0.1, thereby computing the optimized monosaccharide type distribution, *M*_1_, for each token.

The original and optimized monosaccharide type probability distributions, *M*_0_ and *M*_1_, were used to pick which token to change and which monosaccharide to change it to. Two vectors, *P*_0_ and *P*_1_, were generated from *M*_0_ and *M*_1_, giving the probabilities of each token being the current monosaccharide type. These vectors were then used to calculate a discrete probability distribution *M*_*s*_ from which to select the monosaccharide to change, where:

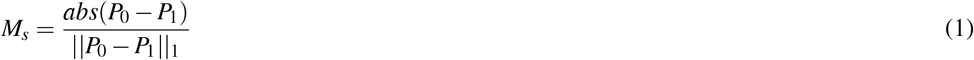

For the *i*^th^ token sampled from *M*_*s*_, *P*_*s*_ is the probability distribution of the *i*^th^ token’s monosaccharide type. To ensure a different glycan was returned after each round of generation, the probability of being the original monosaccharide was changed to 0 in *P*_*s*_, with the altered *P*_*s*_ referred to as 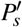 Finally, the new monosaccharide was sampled from 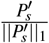 and placed into the glycan structure. To avoid using potentially unrealistic glycosidic bonds while calculating the gradient, once the new monosaccharide was selected, the anomeric conformations and carbon linkages encoding of substituted monosaccharides were left unspecified (using the same “mask” token to mark them as such). Since any context (including glycosidic bond configuration) can impact glycan functionality, it was preferred to calculate the gradient for the next substitution using only verified features of the glycan structures. In future implementations, this might be addressed instead by expanded generative processes to sample updated glycosidic bonds simultaneously, or potentially filtering out glycans unable to be synthesized in an organism of interest [43] depending on the intended use case.

This process was iterated for each starting glycan, providing a series of glycans that generally had increasing probabilities of being immunogenic until a known immunogenic glycan was found or a fixed number of structures (10 in the presented results) were generated (Fig S1D).

The validation set of glycans for the generative process were taken from sets of matching glycan structures (ignoring monosaccharide identities) where each set contained a sufficient number of immunogenic and non-immunogenic pairs. Of the 20 sets of matched glycans structures with at least one immunogenic and one non-immunogenic glycan, 4 were selected for which there were sufficient positive and negative examples (more than 5) and immunogenicity classification was strong (*>* 80% accuracy), thus providing higher confidence in the interpretation of results. From these 4 sets, there were 14 immunogenic and 8 non-immunogenic glycans in total.

## SUPPLEMENTAL

**Table S1.** ABA preferentially binds a glycan with an OS modification on the terminal galactose but does not bind the same glycan when the OS modification is present on C-6. All three glycans are represented in our encoding as GalOS(b1-4)Glcb based on the modification-position agnostic monosaccharide vocabulary found in SugarBase. RFU values are reported from the CFG mammalian microarray v5.0 data used by Coff et al. [14] with ABA at 100 *μ*g available from http://www.functionalglycomics.org:80/glycomics/HFileServlet?operation=downloadRawFile&fileType=DAT&sideMenu=no&objId=1004226. The binding label determined by Coff et al. [14] via the MAD technique is reported in the 4^th^ column.

| CFG v5.0 Probe Num. | Probe | RFU (100 μg ABA) | MAD-determined binding |
| --- | --- | --- | --- |
| 25 | (3S)Galb1-4Glc-SP8 | 1066 | Bound |
| 42 | (6S)Galb1-4Glc-SP0 | 34 | Not bound |
| 43 | (6S)Galb1-4Glc-SP8 | 37 | Not bound |

**Fig S1.**
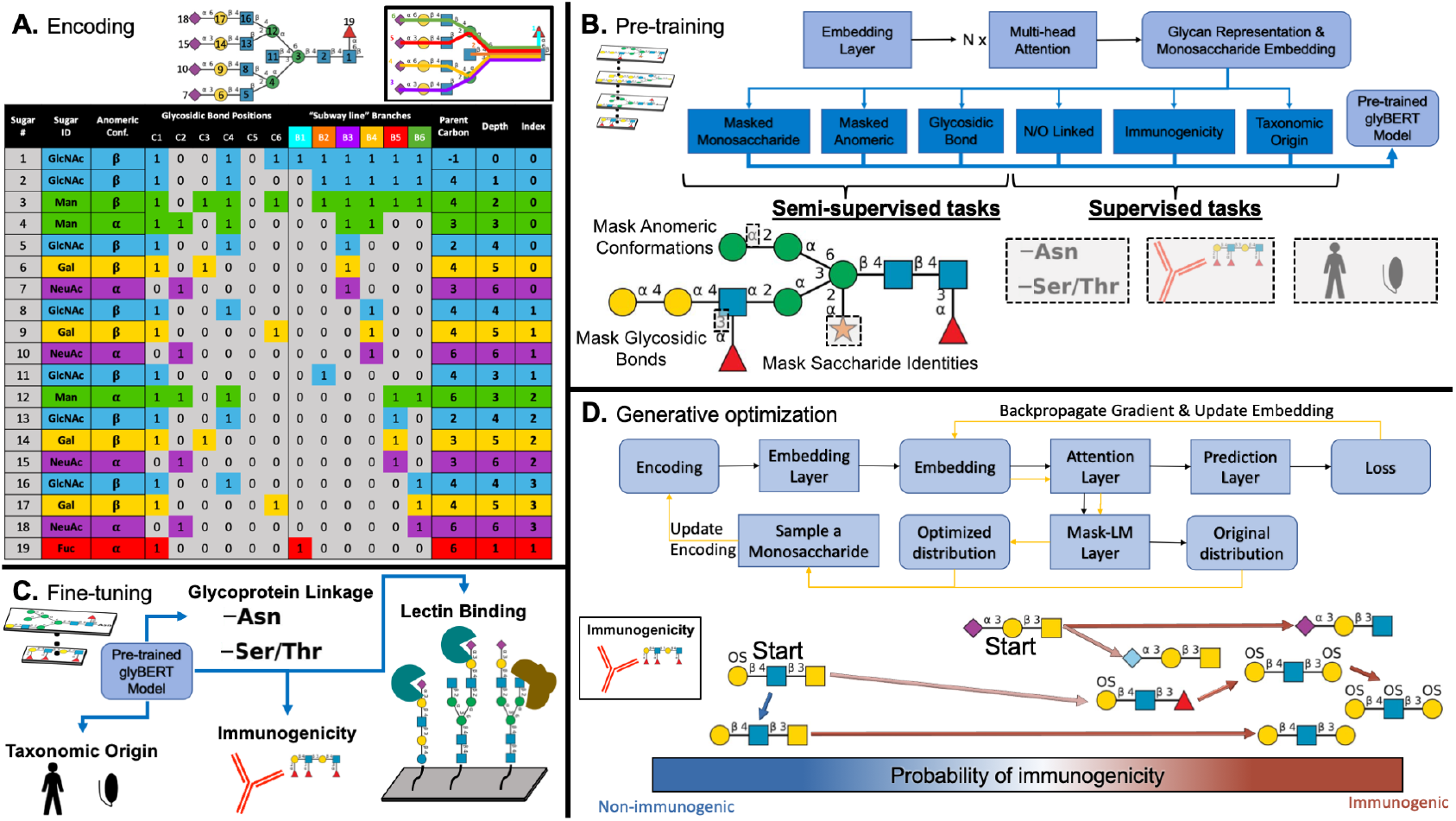
Overview of glycan encoding and glyBERT pre-training, fine-tuning, & generative exploration. **(A)** A simplified example of the glycan encoding used here with fewer bits and without the “START” and “STOP” tokens. Each monosaccharide in a glycan is fully represented in its own row, with an explicit representation of branching using the “subway line” approach demonstrated in the inset in the top right of the panel. The indicator variable columns used to indicate the involvement of carbons in glycosidic bonds or the “subway lines” that pass through each monosaccharide are treated as binary numbers and collapsed into decimal numbers to simplify encoding representations for the model. **(B)** The model architecture and prediction tasks used in pre-training glyBERT. **(C)** Fine-tuning of glyBERT to build specialized glycan representations specific for each prediction task. **(D)** The model architecture used in the generative process. For each generative iteration, there are two components represented by black and yellow arrows. Black arrows indicate the first component, with the goal of getting the status of the current glycan structure before making any changes or updates. From the original encoding, simply passing through the embedding, attention, and prediction layers yield the current monosaccharide embeddings and the current loss towards the optimization target. Yellow arrows indicate the second component, optimization and updating. Embeddings are updated by utilizing the calculated gradient to get new probability distributions for each potential monosaccharide substituent. The final discrete update in the encoding is then generated by random sampling from these probability distributions.

**Fig S2.**
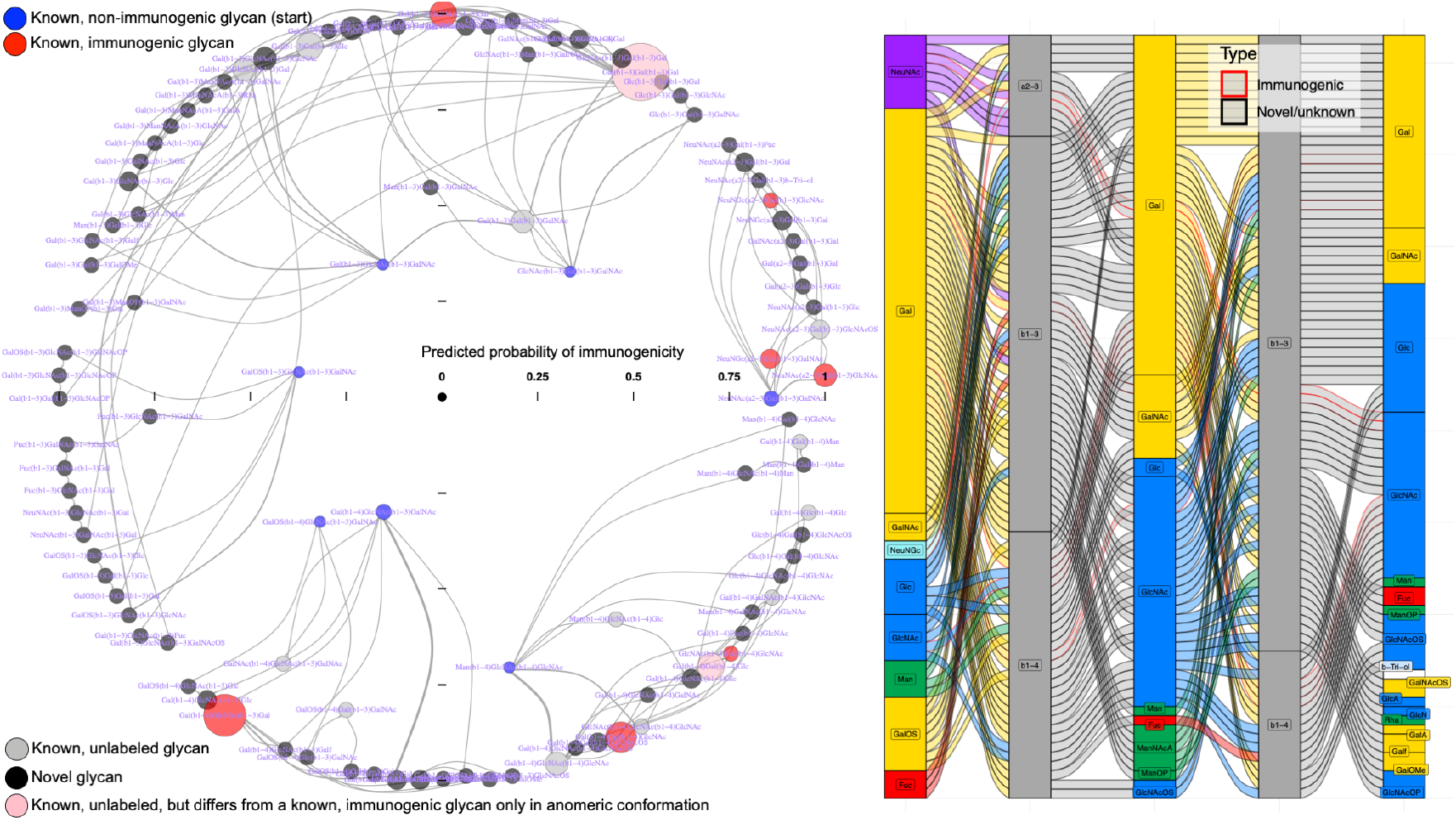
Complete generative exploration of glycans’ structural relationship to immunogenicity. Generative paths from seven non-immunogenic glycans, including the four in Fig 4 along with three additional glycans from which known glycans were discovered which were unlabeled but shared the same carbon attachments and monosaccharide identities as known immunogenic glycans with differing anomeric conformations. In general, the glycans generated here showed similar insights as Fig 4 into the features driving glyBERT’s prediction glycan immunogenicity in this structural context: Gal & Gal derivatives were most common at the non-reducing-end terminal position with GlcNAc or Gal monosaccharides appearing frequently in the middle position, although Gal & GalNAc are better represented here compared to Fig 4. The largest apparent difference from this generative exploration compared to Fig 4 is the presence of glycans with reducing-end terminal GalNAc with high predicted probability of immunogenicity, highlighting other drivers of immunogenicity learned by glyBERT relevant in this structural context.

